# Microfluidic Antibody Affinity Profiling for In-Solution Characterisation of Alloantibody - HLA Interactions in Human Serum

**DOI:** 10.1101/2020.09.14.296442

**Authors:** Matthias M. Schneider, Tom Scheidt, Ashley J. Priddey, Catherine K. Xu, Mengsha Hu, Sean R. A. Devenish, Georg Meisl, Christopher M. Dobson, Vasilis Kosmoliaptsis, Tuomas P. J. Knowles

**Affiliations:** Centre for Misfolding Diseases, Department of Chemistry, University of Cambridge, Lensfield Road, Cambridge CB2 1EW, United Kingdom; Department of Surgery, University of Cambridge, Addenbrooke’s Hospital, Hills Road, Cambridge CB2 0QQ, UK; Fluidic Analytics, Unit A, The Paddocks Business Centre, Cherry Hinton Rd, Cambridge CB1 8DH, UK; NIHR Blood and Transplant Research Unit in Organ Donation and Transplantation, University of Cambridge, Hills Road, Cambridge CB2 0QQ, UK; NIHR Cambridge Biomedical Research Centre, Hills Road, Cambridge CB2 0QQ, UK; Cavendish Laboratory, Department of Physics, University of Cambridge, JJ Thomson Ave, Cambridge CB3 0HE, UK

## Abstract

The detection and characterisation of antibodies in human blood is a key for clinical diagnostics and risk assessment for autoimmunity, infectious diseases and transplantation. Antibody titre derived from immunoassays is a commonly used measure for antibody response, but this metric does not resolve readily the two fundamental properties of antibodies in solution, namely their affinity and concentration. This difficulty originates from the fact that the fundamental parameters describing the binding interaction, affinity and ligand concentration, are convoluted into the titre measurement; moreover, the difficulty of controlling the surface concentration and activity of the immobilised ligand can make it challenging to distinguish between avidity and affinity. To address these challenges, we developed microfluidic antibody affinity profiling, an assay which allows the simultaneous determination of both affinity and antibody concentration, directly in solution, without surface immobilisation or antibody purification. We demonstrate these measurements in the context of alloantibody characterisation in organ transplantation, using complex patient sera, and quantify the concentration and affinity of alloantibodies against donor Human Leukocyte Antigens (HLA), an extensively used clinical biomarker to access the risk of allograft rejection. These results outline a path towards detection and in depth profiling of antibody response in patient sera.

## Introduction

Non-covalent protein-protein interactions underlie many biological and physiological processes, including protein self-assembly,^1–3^ protein aggregation,^4–6^ antibody-antigen recognition,^7–9^ muscle contraction,^10^ and cellular communication.^11^ One fundamental application of measuring protein-protein interactions is based on immuno-assays for the detection of biomarkers in body fluids, primarily human serum, related to various diseases including cancer,^12–15^ protein misfolding diseases,^16–21^ auto-immune diseases,^22^ and graft rejection.^23^ In the latter, detection of antibodies against donor Human Leukocyte Antigens (HLA), termed alloantibodies, in patient serum serve as strong indicator for potential rejection of transplants. The analysis and characterisation of these biomarkers is essential for pre-transplant assessment and post-transplant immune monitoring.^24–27^

Current approaches for the detection and characterisation of antibodies, including alloantibodies, rely mostly on surface immobilisation of one of the binding partners, such as in enzyme-linked immunosorbant assays (ELISA),^28–30^ bead-based multiplex assays,^31–34^ and surface plasmon resonance (SPR) spectroscopy.^35–37^ The requirement for protein immobilisation is associated with a number of challenges, such as non-specific interactions with the surface or suppressed accessibility due to alterations of substrate and ligand structures.^38–40^ Furthermore, avidity effects caused by dense substrate occupation are hard to control in surface based measurements.^41^ Additionally, the hook/prozone effect, most prevalent in sandwich immunoassays or complement interference, can result in false negative measurements.^42–44^ More generally, for measurements on surfaces, the fundamental parameters of antibodies in solution, namely their affinity and concentration, are challenging to resolve due to the difficulty in controlling the concentration of the surface-bound species. For instance, the commonly used EC50 value obtained in surface measurements is only weakly dependent on the affinity of the interaction for strong antibody binding, as illustrated in Fig.1a.

**Figure 1:**
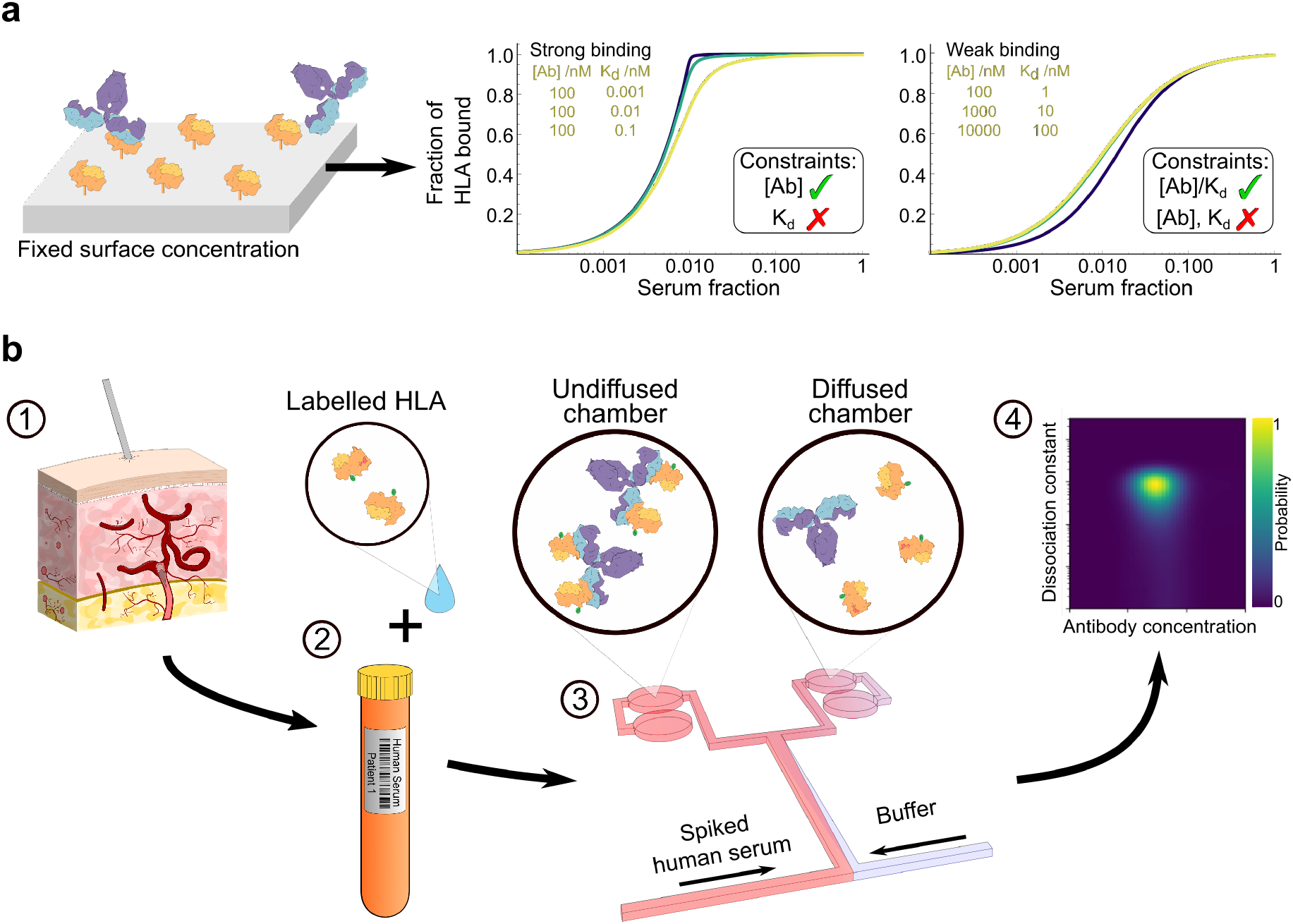
(**a**) Basic Principle of surface based antibody-binding measurements. In a strong binding regime, the antibody concentration can be determined, in a weak binding regime the ratio between antibody concentration and dissociation constant, *K*_*d*_, can be determined only. (**b**) Basic principle of applying MDS for clinical samples. (1) Patient-derived human serum (2) is incubated with different concentrations of labelled HLA to allow binding. (3) The effective size of the complex is determined by microfluidic diffusional sizing, from which (4) the dissociation constant *K*_*d*_ and the antibody concentration [*Ab*] are determined. As it is an in-solution approach, both *K*_*d*_ and antibody concentration become accessible. The posterior shows the probability distribution, whereby yellow stands for high, blue for low probability that the according parameters *K*_*d*_ and [*Ab*] are found at the respective values.

To overcome the limitations of surface-based immunoassays, we demonstrate here an in-solution Microfluidic Antibody Affinity Profiling (MAAP) apporach, allowing quantitative measurement of biophysical parameters governing specific antibody-antigen interactions in unpurified human serum and its complex background with more than 10 million different proteins.^45^ Unlike existing microfluidic immunoassays, many of which rely on surface immobilisation,^46,47^ the approach described in this paper operates fully in solution. We use a strategy based on microfluidic diffusional sizing, which tracks the spatial and temporal evolution of a fluorescently labelled protein in a microfluidic channel under laminar flow conditions to determine its hydrodynamic radius, *R_h_* (Fig. 1) and hence effective molecular weight. When an antigen molecule interacts with an antibody in solution, its effective molecular weight increases to that of the antibody-antigen complex and hence its diffusion coefficient decreases.

We focus here on humoral responses against HLA, also known as the human major histocompatibility complex (MHC), a biomolecule of key clinical significance. The extensive polymorphism of the HLA system, evolved to enable immune protection against an ever-changing environment of human pathogens, is a major barrier in organ and cell transplantation.^48^ Exposure to donor HLA through pregnancy, transfusion and/or transplantation leads to development of alloantibodies, principal mediators of acute and chronic allograft loss.^23,49^ Detection and characterisation of alloantibodies is essential for evaluation of donor-recipient compatibility and to facilitate post-transplant immune monitoring and provide individual therapeutic intervention.^50–52^ In current clinical practice, this is mainly performed using solid phase assays which suffer from the aforementioned disadvantages of surface-based techniques. Such semi-quantitative approaches do not allow the full characterisation of fundamental biophysical properties of the humoral alloresponse such as alloantibody levels (concentration) and the affinity of alloantibody-HLA interaction.^27,53–55^ In the following, we show that MAAP is an advanced technique capable of quantifying an analyte in human serum under native solution conditions to yield physiologically relevant results and thus propose this platform as an additional procedure for immuno-profiling in human serum.

## Results and Discussion

### Binding Interactions of Covalently Labelled HLA

In order to detect alloantibodies in solution using HLA, a microfluidic diffusional sizing (MDS) platform was used that enables determination of binding parameters by measuring the hydrodynamic radius, *R_h_*, of a fluorescently labelled protein, as previously described.^56–59^ We used a reliable and stable labelling strategy for fluorescence detection, relying on NHS-Chemistry (Fig. S1a) for N-terminal labelling, which yields highly pure HLA with labelling stoichiometry of 0.33 to 1.55, depending on the variant (Fig. S1c-f). This allows control of the stoichiometry of the binding interaction more accurately than traditional strategies utilising biotin-streptavidin-HLA complexes which are highly heterogeneous (Fig. S1b). A fluorophore in the far-red spectral region was chosen (*λ*_em,max_ = 650 nm), since serum autofluorescence is minimised in this spectral region (Fig. S2e).

Rapid and physiologically accurate investigation of antibodies in human samples is crucial for clinical evaluation. Therefore, immunoassays must be able to cope with untreated samples and not be influenced by non-specific binding to surfaces, by the hook/prozone-effect, and by surface-mediated avidity effects.^42–44^ To validate the applicability of the assay for quantification of alloantibody-HLA interactions, the binding of the HLA variant A*03:01 to the mouse derived anti-human monoclonal antibody W6/32, which specifically recognises a monomorphic epitope on all HLA class I molecules that includes both the heavy chain and the β_2_-microglobulin chain,^60^ was investigated first in pure buffer. The hydrodynamic radius, *R_h_*, of pure HLA A*03:01 was determined to be 3.47 *±* 0.13 nm (Fig. 2a), which is consistent with the expected radius for a natively folded protein with a molecular weight of approximately 55 kDa (Fig. 2b). Upon addition of a 380-fold excess of antibody W6/32, a significant increase of the hydrodynamic radius to *R_h_* = 5.01 *±* 0.13 nm was observed, indicating an interaction between both species.

**Figure 2:**
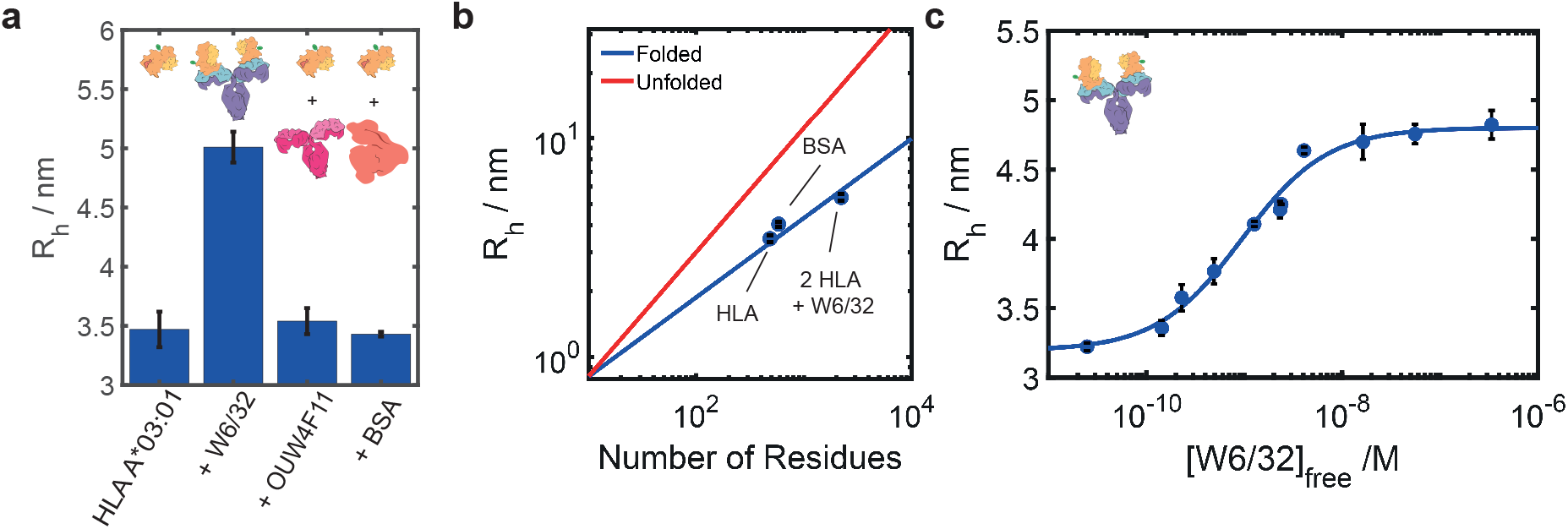
(**a**) Binding experiments of HLA A*03:01 using diffusional sizing. Bars show the average of triplicate measurements with the error bars representing the standard deviation. A significant change in hydrodynamic radius indicates binding of 5 nM HLA A*03:01 to 1.9 *μ*M W6/32. In contrast, no binding can be observed between HLA and OUW4F11 or BSA. (**b**) Correlation of hydrodynamic radii with the number of residues.^61,62^ The radii determined for HLA both free and bound to the antibody W6/32 agree well with the assumption of a folded protein. (**c**) Binding curve of 476 pM HLA A*03:01 with varying concentration of antibody W6/32. The blue points give the hydrodynamic radius of each equilibrated sample, averaged over the data of at least three replicates, and the blue curve is the best fit (see Materials and Methods for details). From the fit, the dissociation constant *K*_*d*_ = 0.7 [0.3, 1.6] nM (95% confidence intervals given in square brackets) could be determined with a ratio of two antigens per antibody.

The hydrodynamic radius of 5.01 nm suggests that two antigens are bound per antibody, which is expected for a bivalent IgG antibody which has two binding sites, and is consistent with the expected size for a natively folded protein complex of 260 kDa (Fig. 2b). Negative control experiments, including alloantibody against fluorescently labelled BSA (Fig. S4a), OUW4F11 (an alloantibody specific to HLA B*08:01) against HLA A*03:01 (Fig. 2a), and human IgG binding against HLA A*03:01 (Fig. S4b), did not show an increase in *R_h_* and, therefore, confirmed that the complex formation is based only on specific interactions.

We next explored whether this approach could yield both the dissociation constant of the interaction and the concentration of the antibody. To this effect, an equilibrium binding curve for the interaction between HLA A*03:01 and antibody W6/32 was measured, yielding a *K*_*d*_ = 0.7 [0.3, 1.6] nM (95% confidence intervals from Bayesian inference given in square brackets) and consistent with a binding ratio of 1 to 2, i.e. a stoichiometry of 2 antigens per antibody (Fig. 2c and Fig. S5). Both cooperative and non-cooperative binding of antibodies have previously been described.^63^ Thus, cooperativity was tested by a Hill plot (Fig. S5),^64^ yielding a Hill parameter *h* = 1.01 *±* 0.15. This shows that the binding of the HLA and the antibody is non-cooperative, ergo binding events occur independently.

### Characterisation of Alloantibody-HLA Interactions in Human Serum

We next determined the applicability of MAAP for the absolute quantification and characterisation of alloantibodies in human serum. We assayed two well-characterised, human monoclonal antibodies, SN23OG6 and OUW4F11, which specifically recognise HLA A*02:01 and HLA B*08:01, respectively,^65^ both in human serum of non-transfused, healthy donors, which did not contain HLA-specific antibodies, and in buffer (PBS). During data processing, we corrected for a weak autofluorescence background signal from both the diffused and undiffused measurement channels. Serum autofluorescence has been reported for human serum in the spectral region of interest (Fig. S2)^66^ and is likely to arise from natural aromatic compounds including haem complexes, found in haemoglobin or bilirubin, which are stabilised by human serum albumin.^67^

As shown in Fig. 3, the hydrodynamic radii of pure HLA obtained in human serum (*R_h_* = 3.27 *±* 0.11 nm for HLA A*02:01 and *R_h_* = 3.22 *±* 0.19 nm for HLA B*08:01), were found to be consistent with theoretical values for natively folded 50 kDa proteins, as well as with the radii obtained in buffer (*R_h_* = 3.22 *±* 0.10 nm for HLA A*02:01 and *R_h_* = 3.16 *±* 0.07 nm for HLA B*08:01), demonstrating the applicability of MDS for measurements in human serum. Importantly, this also shows that measurement in human serum did not affect the antigen size compared to the measurements in PBS, suggesting presence of serum proteins in the sample do not affect the measurements.

**Figure 3:**
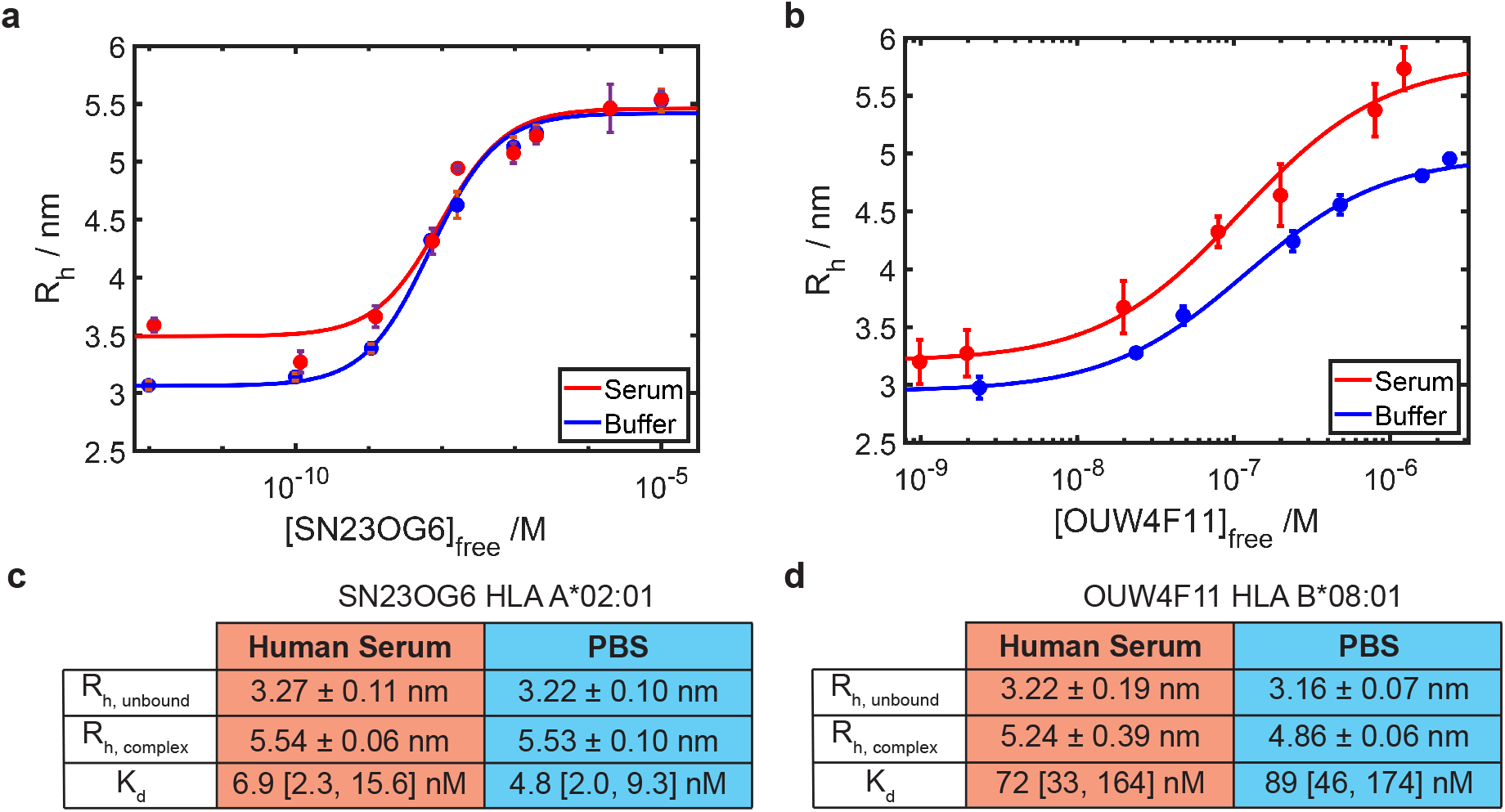
(**a**) Binding curve of 5 nM HLA A*02:01 against antibody SN23OG6 in human serum (red) and PBS (blue). The measurements in serum and PBS are in good agreement with each other, yielding *K*_*d*_ values of 6.9 [2.3, 15.6] nM and 4.8 [2.0, 9.3] nM, respectively, with a binding ratio of 2 antigens per antibody. (**b**) Binding curve of 1.2 nM HLA B*08:01 against antibody OUW4F11 in human serum (red) and PBS (blue). Again, the *K*_*d*_ = 72.1 [32.9, 163.6] nM in serum and *K*_*d*_ = 89.1 [45.7, 173.8] nM in PBS with a binding ratio of 1 to 2 are in good agreement. Summary of the hydrodynamic radii for the fully unbound antigens and of the dissociation constants in human serum in comparison to their values in pure buffer, demonstrating consistent values under both conditions (**c**) for SN23OG6 against HLA A*02:01 and (**d**) for OUW4F11 against HLA B*08:01 (Fig. S6).

Assessment of these two alloantibody-HLA interactions showed that microfluidic affinity measurements were independent of the buffer conditions used (Fig. 3a-b). The interaction between SN23OG6 against HLA A*02:01 yielded a dissociation constant of *K*_*d*_ = 6.9 [2.3, 15.6] nM in human serum and was in good agreement with *K*_*d*_ = 4.8 [2.0, 9.3] nM in buffer. Similarly, for the interaction of antibody OUW4F11 against HLA B*08:01, the *K*_*d*_ = 72.1 [32.9, 163.6] nM in human serum which was consistent with *K*_*d*_ = 89.1 [45.7, 173.8] nM determined in buffer. The different saturation levels between the two media conditions are within the CI and most likely reflect minor conformational variations between individual antigens. Analysis of the above alloantibody-HLA interactions in PBS using biolayer interferometry showed dissociation constants in a similar order (*K*_*d*_ = 5.6 *±* 0.03 nM for SN23OG6 vs. HLA A*02:01 and *K*_*d*_ = 314.9 *±* 0.04 nM for OUW4F11 against HLA B*08:01, Fig. S7).

Taken together, these data show the general feasibility of binding measurements in complex media with diffusional sizing, with hydrodynamic radii, affinities and stoichiometric parameters consistent with theoretical values and measurements under unperturbed (buffer) conditions. The results show that the binding of HLA, even in the background of a complex solution such as human serum, is solely based on specific interactions and not influenced by any other protein species that are abundant in human serum including soluble HLA and β_2_microglobulin.^68^ More generally, these results suggest that MAAP can be used as a platform for molecular level characterisation of protein-protein interactions in complex mixtures such as body fluids.

### Quantification of Alloantibody-HLA Interactions in Human Serum

Simultaneous determination of both affinity and alloantibody concentration in patient samples is a key advantage of our method compared to traditional assays. Through varying the concentration of both labelled (i.e. HLA) and unlabelled (i.e. alloantibodies in human serum) species, it becomes possible to properly constrain the probability distribution of unknown parameters for the interaction (*K*_*d*_ and antibody concentration) as demonstrated by Bayesian inference analysis (see Methods). This is the key advance of this technology as compared to similar assays previously used to describe affinity measurements. In order to verify the robustness of our method for determination of absolute parameters for reactive antibody species, we spiked human serum from non-sensitised donors with HLA-specific monoclonal antibody in a blinded manner, i.e. the final antibody serum concentrations were not revealed to the person performing the analysis. The interaction investigated was that between alloantibody SN23OG6 and HLA A*02:01 (Fig. 4). For antibody concentrations of 1 nM or above, we were able to determine both the dissociation constant, *K*_*d*_, and the concentration of specific antibody, [Ab]_spec_, in doped serum simultaneously. The *K*_*d*_ determined in all cases was consistent with previous results for the interaction (as shown above), and the concentration determined was in good agreement with the expected antibody concentration (Fig. 4). This was the case even when the concentration of antibody binding sites was approximately equal to the dissociation constant, *K*_*d*_ (Fig. 4a), demonstrating that our method enables accurate determination of antibody concentrations around the *K*_*d*_.

**Figure 4:**
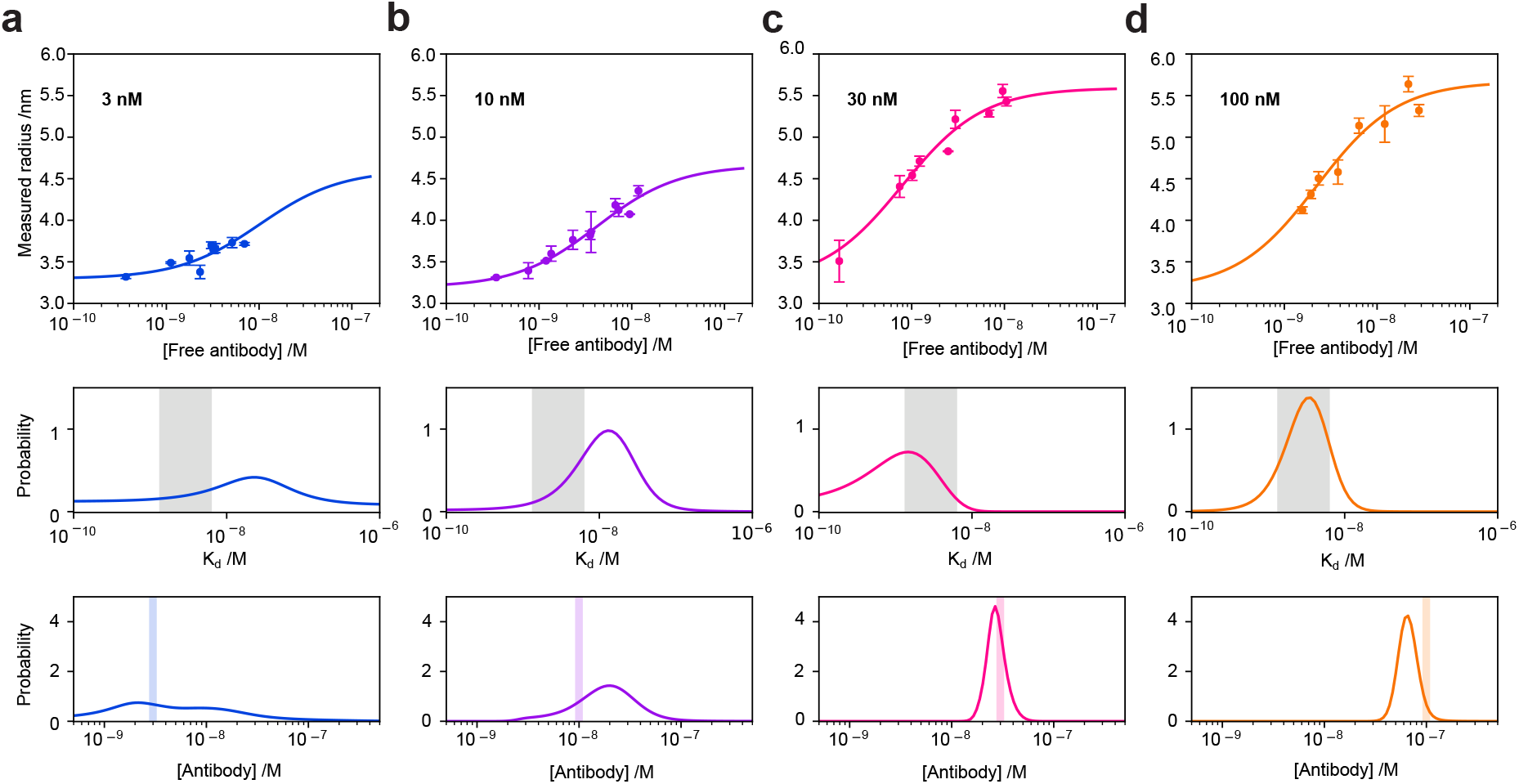
Binding curves (top row) of measured data and best fits for the interaction of A*02:01 against SN23OG6 antibody spiked into alloantibody-negative human serum at concentrations (**a**) 3 nM, (**b**) 10 nM, (**c**) 30 nM and (**d**) 100 nM. The analyses were performed assuming that both the antibody concentration and affinity were unknown. In all four cases, the Kd determined through MAAP is consistent with the *K*_*d*_ obtained through fitting all datasets combined, using the known concentrations; probability distributions over *K*_*d*_ resulting from each dataset are shown alongside the 95% confidence intervals (grey) obtained by considering all data (middle row). The experimentally determined antibody concentrations are also in good agreement with the known concentration (bottom row), assuming a binding stoichiometry of 1:2 antibody:HLA. Probability distributions over antibody concentration are overlaid with the experimental error range (shaded region) for the known antibody concentrations.

### Quantification of Alloantibody-HLA Interactions in Patient Sera

We next investigated the ability of our immunoassay to quantify HLA-specific antibodies in the serum of a kidney transplant patient. The patient became sensitised after transplantation with a kidney allograft expressing the HLA A*24:02 alloantigen. Analysis of post-transplant sera using the Luminex single antigen bead assay (standard of care in the clinical setting) showed a complex profile with reactivity against the priming antigen (A*24:02), cross-reactivity against A*02:01 (A*24:02 and A*02:01 are part of a common serological cross-reactive HLA epitope group), and additional reactivity to A*01:01 (no known serological cross-reactivity to A*24:02 and A*02:01^69^). The mean fluorescence intensity values detected by Luminex were 19314 a.u. for A*24:02, 18653 a.u. for A*02:01 and 7810 a.u. for A*01:01 (Fig. 5a). As shown in Fig. 5b, for the serum interaction with HLA A*02:01, we determined a concentration of alloantibody of [Ab]_spec_ = 3.0 [1.2, 5.8] nM and a *K*_*d*_ = 0.13 [0.01, 1.38] nM, assuming a binding ratio of 1 to 2. Similarly, for the same patient serum interaction with A*24:02, we detected an antibody concentration of, [Ab]_spec_ = 19.3 [10.8, 49.3] nM and an affinity of *K*_*d*_ = 1.3 [0.2, 9.7] nM. Thus, we were able to deconvolute the fundamental biophysical properties (affinity and alloantibody concentration) of the humoral response in a complex patient serum and differentiate the reactivity against the priming alloantigen (A*24:02) and a cross-reactive alloantigen (A*02:01) demonstrating higher antibody concentration against the priming HLA. Importantly, we could not demonstrate an interaction between HLA A*01:01 and the patient serum, despite a relatively high MFI value of 7810 a.u. from the Luminex assay. Output from the Luminex assay is avidity driven and is considered semi-quantitative;^40^ accordingly, an interaction at the MFI level showed here against A*01:01 would be considered as clinically significant (e.g. potential donors expressing A*01:01 would typically be excluded for a patient with similar levels of reactivity on Luminex). Taken together, the data highlight the potential of MAAP to provide immunologically relevant information not attainable by currently available techniques and to quantify antibody interactions against proteins that share highly similar structures, such as the HLA system.

**Figure 5:**
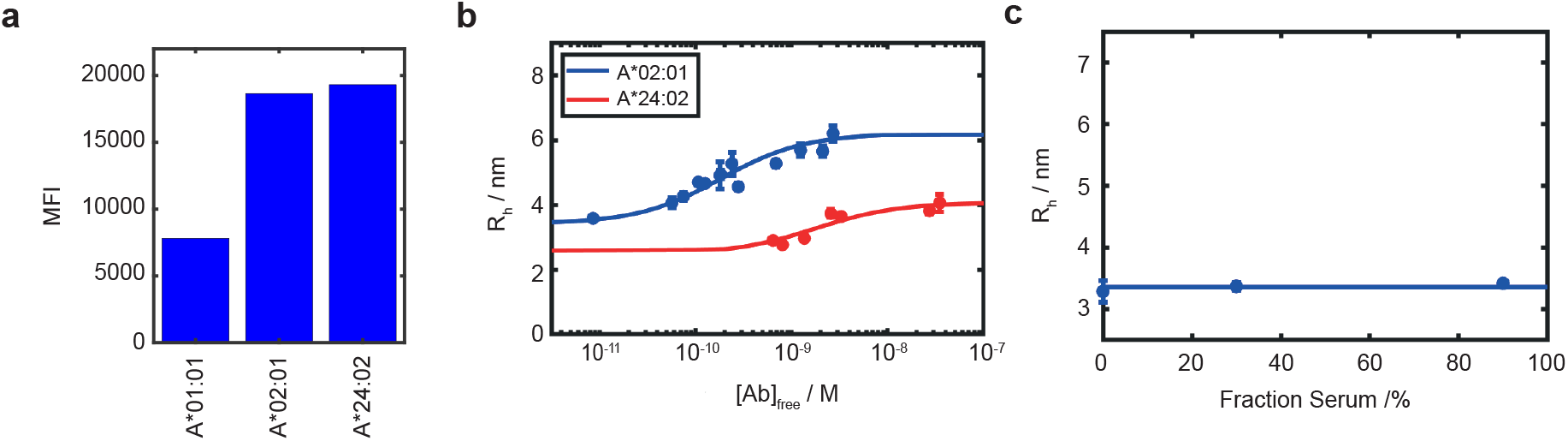
Quantification of reactivity against different HLA variants in serum from a transplant patient. Antibody binding profile as detected on Luminex Single antigen beads for HLA A*01:01, HLA A*02:01 and HLA A*24:02 (MFI: Mean Fluorescence Intensity). (**b**) Binding Curves for the interaction of alloantibodies in patient serum against HLA A*02:01 and A*24:02, relating the hydrodynamic radius, *R_h_*, to the measured concentration of antibody. The individual data points represent the measured data, the solid line the fit. The error bars report the standard deviation for triplicate measurements. For HLA A*02:01 isoform, an antibody concentration [Ab]_spec_ = 3.0[1.2, 5.8] nM and a dissociation constant *K*_*d*_ = 0.13 [−, 1.38] nM were determined. For HLA A*24:02 isoform, we determined the *K*_*d*_ = 1.3 [0.2, 9.7] nM and [Ab]_spec_19.3 = [10.8, 49.3] nM. (**c**) Titration Curve for the same patient serum against 1 nM HLA A*01:01, with different serum concentration. As can be seen, no binding is detected.

## Conclusion

In this study, we have shown for the first time that, using an in-solution technique, namely Microfluidic Antibody Affinity Profiling (MAAP), it is possible to determine both dissociation constant, *K*_*d*_, and the absolute concentration of antibody binding sites, which may manifest as the binding stoichiometry in samples of known antibody concentrations, or the total antibody concentration in unknown samples, through a measurement of effective hydrodynamic radii at different antibody and antigen concentrations. By applying this platform to measure immunologically-relevant interactions between specific antibodies and HLA, we were able to determine quantitative biophysical parameters describing the binding event fully, even in such a complex medium as human serum. The determined dissociation constants range between 10^−10^ M and 10^−8^ M and are consistent with previous work.^70^ However, our in-solution approach avoids the commonly reported disadvantages of surface-based assays and does not require serum preparation to reduce nonspecific binding thereby enabling determination of fundamental parameters of humoral responses under physiological conditions.

Our results suggest applicability of this method in a wide range of investigations aiming to understand the role of both abundance and dissociation constants implicated in clinically relevant immune responses. For example, further insights into the complex process of graft rejection may be obtained through investigations of the correlation of the concentration and affinity to the occurrence of an immune response. Finally, our results indicate that the platform can be of general use for diagnostics beyond histocompatibility testing, such as immuno-profiling in auto-immunity, in infectious diseases, and to monitor immune responses after vaccination, or the detection of biomarker levels for various diseases in human serum.^15,16^

## Acknowledgement

The research presented in this manuscript has received funding from the European Research Council under the European Union’s Seventh Framework Programme (FP7/2007-2013) through the ERC grant PhysProt (agreement no. 337969) and under European Union’s Horizon 2020 research and innovation programme (ETN grant 674979-NANOTRANS). We gratefully acknowledge financial support from the Engineering and Physical Sciences Research Council (EPSRC) and Frances and Augustus Newman Foundation. We also acknowledge funding from the NIHR Blood and Transplant Research Unit in Organ Donation and Transplantation at the University of Cambridge and from an NIHR Fellowship (PDF-2016-09-065, VK). The views expressed are those of the authors and not necessarily those of the National Health Service, the National Institute for Health Research, the Department of Health, or National Health Service Blood and Transplant. We are grateful to Dr Arend Mulder and Prof Frans Claas for provision of monoclonal antibodies SN23OG6 and OUW4F11. Ethical Approval for work involving human serum was granted from NRES Committee North-East York, IRAS number 167211.

## Conflict of Interest

S.R.A.D. is an employee of Fluidic Analytics, which is developing and commercializing microfluidic diffusional sizing instrumentation including the Fluidity One-W Serum used here. V.K. is a consultant with Fluidic Analytics Ltd. T.P.J.K. is a member of the board of directors of Fluidic Analytics Ltd.

## Supporting Information

The following files are available free of charge online.

## Notes

### Competing Interest Statement

Sean R.A. Devenish is an employee of Fluidic Analytics, which is developing and commercializing microfluidic diffusional sizing instrumentation including the Fluidity One-W Serum used here.
Vasilis Kosmoliaptsis is a Consultant with Fluidic Analytics Ltd. Tuomas P.J.Knowles is a member of the board of
directors of Fluidic Analytics Ltd.

