## Supporting Information for "Microfluidic Antibody Affinity Profiling for In-Solution Characterisation of Alloantibody - HLA Interactions in Human Serum"

### Supplementary Information

### Materials and Methods

#### Materials

HLA monomers were obtained through the NIH Tetramer Core Facility in Emory, Atlanta, US, in PBS. PBS and Alexa Fluor 647 was purchased from Thermo Fisher Scientific Inc., Waltham, US. Human IgG ab205198 was from Abcam, Cambridge, UK. All other chemicals were from Sigma Aldrich. All PBS was supplemented with $NaN_3$ (0.02 % (w/v)).

Plate reader experiments were performed on a Clariostar BMG Labtech, Ortenberg, DE instrument. Size exclusion chromatography was performed on a Superdex 200 increase 10/300 gL column (GE Healthcare, Chicago, US) on an AKTA Pure protein purification system (GE Healthcare, Chicago, US). All microfluidic experiments were performed on a Fluidity One-W Serum instrument(Fluidic Analytics, Cambridge, UK). The basic principle of MDS has been described before.[1] In brief, labelled protein and auxiliary buffer are introduced alongside one another at the beginning of an extended diffusion chamber. Due to the small channel size, laminar flow can be assumed, meaning that the particles can move into the buffer stream by diffusion only, whereby the rate depends on the size of the molecular complex (Fig. 1). At the end of the diffusion chamber, the stream is split and the fluorescence of both the diffused and the undiffused material is measured. From the ratio between the fluorescence in both chambers, the hydrodynamic radius, $R_h$, of the protein can be determined.

#### Determination of Autofluorescence in Human Serum

Human serum from non-sensitised volunteers (not containing HLA-specific antibodies) was supplemented with PBS and the fluorophore Alexa Fluor$^{TM}$ 647 to yield fluorophore concentrations between 10 pM and 1 μM in serum. Similar dilutions of fluorophore in buffer were prepared for comparison. Subsequently, both absorption spectra (Fig. S2 a-b) and the emission spectra upon excitation at two wavelengths $\lambda_{ex,1}$ = 481 nm and $\lambda_{ex,2}$ = 632 nm (Fig. S2 c-d)were recorded on a plate reader.

#### Labelling of HLA with Alexa Fluor 647 fluorophore

To HLA (typically 1 nmol, 1 equiv.) in 0.1 M $NaHCO_3$ (pH = 8), Alexa Fluor 647 *N*-hydroxysuccinimide ester (in DMSO, 3 equiv.) was added. The reaction mixture was incubated for 1 h at ca. 20 °C, protected from light. The sample was purified by size exclusion chromatography with a flow rate of 0.5 mL/min and PBS as eluent buffer, to yield labelled HLA (typically around 0.8 nmol, DOL between 0.33 and 1.55).

#### MAAP measurements in PBS

Labelled HLA, together with a varying concentration of the antibody of interest, were added and diluted in PBS (supplemented with 0.02 % Tween-20). The samples were incubated at room temperature for approximately 30 minutes. Subsequently, the size was determined by MDS. The same protocol was followed for negative controls with fluorescently labelled BSA.

#### Binding Experiments in Human Serum

All binding experiments in human serum were carried out using human serum from non-sensitised volunteers (not containng HLA-specific antibodies) or from a kidney transplant patient. For all binding experiments in human serum, HLA (typically at a total concentration of 5 nM) was incubated with a varying concentration of specific antibodies or serum fractions at room temperature for 30 minutes in human serum and measured by MAAP. When fitting the data, the background fluorescence intensity of both the diffused and the undiffused channel was subtracted from the sample values at the relevant serum concentration.

#### Bayesian Analysis

The dissociation constant, $K_d = \frac{[Ab][H]}{[AbH]}$, and where necessary the antibody binding site concentration, $[Ab]_0$, were determined through Bayesian inference. The hydrodynamic radii were measured based on the amount of protein that diffuses into the distal channel; in order to relate the measurements, we introduce the parameters $\rho_f$ and $\rho_b$ as the fractions of free and antibody-bound HLA, respectively, that diffuse into the distal channel. We can therefore express the fraction of HLA that diffuses into the distal channel, $f_d$, as:

$$f_d = \frac{([AbH]\rho_b + ([H]_0 - [AbH])\rho_f)}{[H]_0} \tag{1}$$

where $[AbH]$ denotes the equilibrium concentration of antibody-HLA complex, and $[H]_0$ the total HLA concentration. Considering mass-balance, we express $[AbH]$ as:

$$[AbH] = \frac{[Ab]_0 + [H]_0 + K_d - \sqrt{([Ab]_0 + [H]_0 + K_d)^2 - 4[Ab]_0[H]_0}}{2} \tag{2}$$

In the experiments where the total antibody concentration was unknown, we substitute $[Ab]_0$ for $\alpha[Ab]_{tot}$, where $\alpha$ is the fraction of serum used in the measurement, and $[Ab]_{tot}$ is the total concentration of antibody binding sites in the stock solution. Assuming a binding stoichiometry of 1:2 antibody:HLA, we therefore obtain the antibody concentration, $\frac{[Ab]_{tot}}{2}$. All confidence intervals used were 95% confidence intervals, corresponding to the standard derivation.

The priors used for $\rho_f$ and $\rho_b$ were flat in linear space, while flat priors in logarithmic space were used for $K_d$ and antibody concentration, where applicable, where employed.

#### Luminex Single Antigen Beads

HLA-specific antibody reactivity in the patient serum was detected using LabScreen single antigen HLA class I detection beads (One Lambda, Canoga Park, CA), as previously described[2]. HLA single antigen bead-defined Ab reactivity was determined using a mean fluorescence intensity (MFI) cut-off threshold of 2000 (MFI cut-off level used clinically in our center and elsewhere to define a positive alloantibody response to a given HLA).

##### Bio-Layer Interferometry

Monoclonal antibody affinity of binding to HLA was determined by bio-layer interferometry (BLI) using the Octet RED96 system (ForteBio, Fremont, California). Antibody was immobilised to anit-human IgG Fc kinetic biosensors. To determine the association phase, sensors were dipped into wells containing soluble, recombinant HLA in a 2-fold titration for 300 seconds so an equilibrium was reached. Next, sensors were placed into buffer alone-containing wells for further 1000 seconds to determine the dissociation phase. Affinity values ($K_d$) were calculated via steady-state analysis as the ratio of on- and off-rate constants ($\frac{k_{off}}{k_{on}}$. All experiments were carried out using standard kinetic buffer (PBS, 0.1% (w/v) bovine serum albumin, 0.02% Tween-20), at a temperature of 30°C and a constant plate shake speed of 1000 rpm.

#### Author Contribution

T.P.J.K., V.K., C.M.D., T.S., M.M.S. and A.J.P. designed the study. M.M.S., T.S., M.H., A.J.P. performed the experiments. A.J.P., S.R.A.D., V.K. and T.P.J.K. provided material. C.K.X., G.M., M.M.S., T.S., V.K. and T.P.J.K. analysed the data. M.M.S., T.S., C.K.X., G.M., V.K. and T.P.J.K. wrote the paper. All authors discussed the results and commented on the manuscript.

### Supplementary Figures

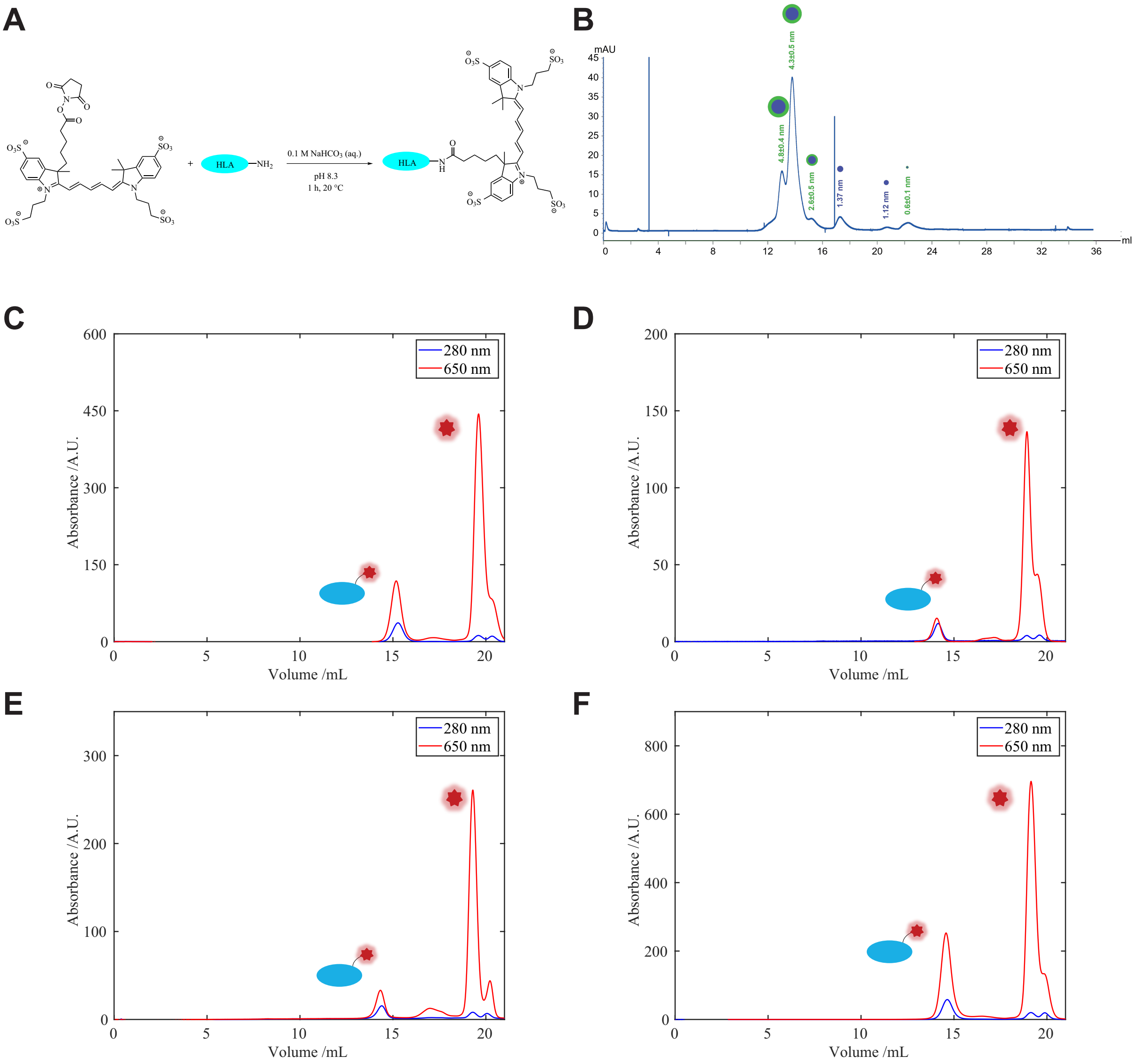


**Figure 1:** Strategy of Covalent labelling. (**A**) Reaction Mechanism of Linking an amine on a protein to Alexa Fluor 647, using an amide coupling. (**B**) Chromatogram for the purification of the streptavidin-HLA complex mixture. The blue circles represent different sizes of different streptavidin-HLA complexes; the green borders represent the Alexa Fluor$^{TM}$ 488 label used in the mixture. (**C**) Chromatogram of the purification of Alexa Fluor 647 labelled HLA A*02:01 (degree of labelling (DOL) 1.22), (**D**) HLA A*03:01 (DOL 0.33), (**E**) HLA B*24:02 (DOL 0.96) and (**F**) HLA B*08:01 (DOL 1.55) after labelling, showing the elution of HLA (blue cartoon with red fluorophore) in one fraction around 14 mL and the excess fluorophore (red star) around 20 mL. The degree of labelling refers to the number of fluorophore per protein molecule.

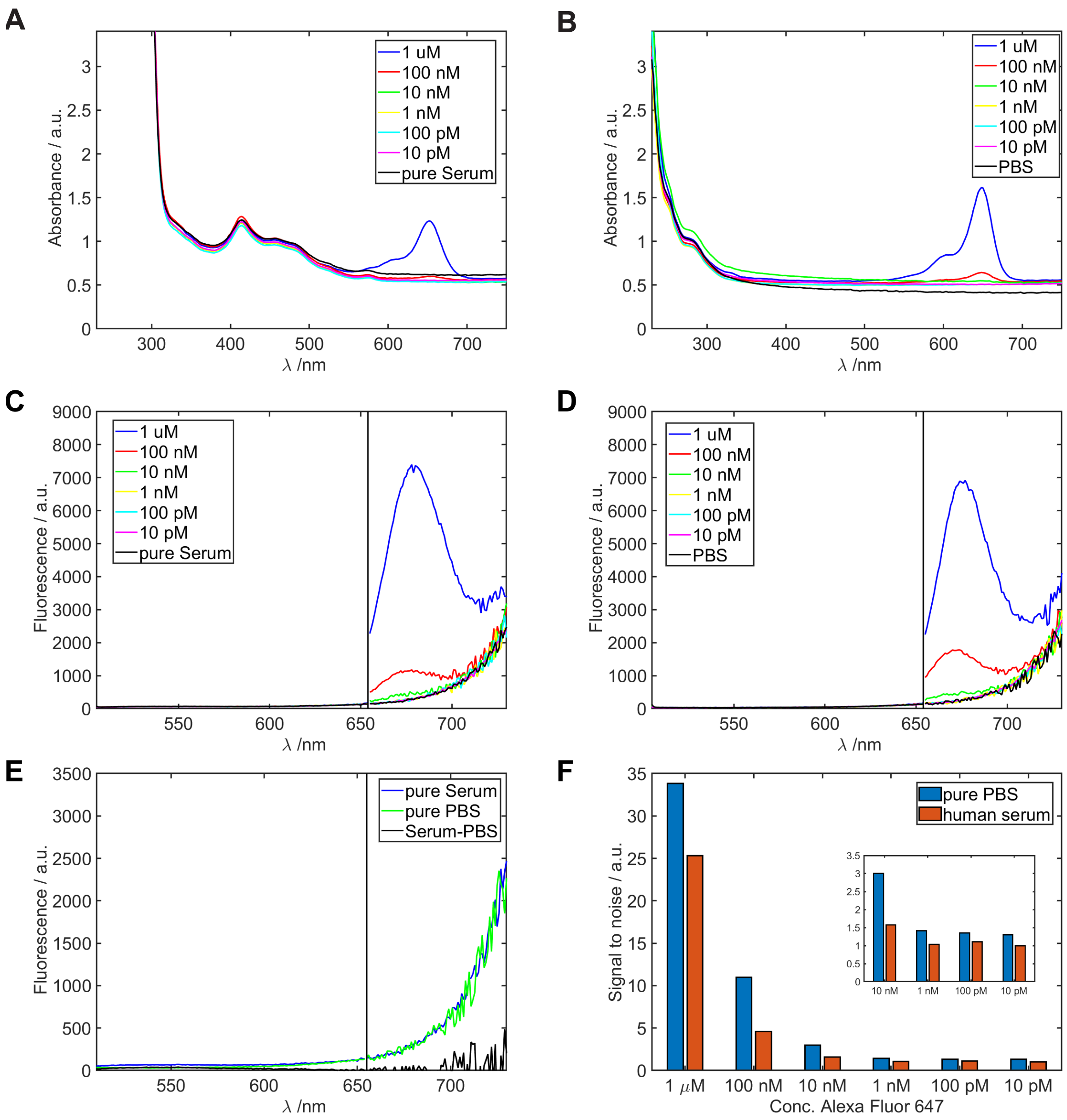


**Figure 2:** (**A**) Determination of the absorbance of varying concentration of Alexa Fluor 647 in human serum and (**B**) in buffer, showing no background absorbance around 663 nm in human serum. (**C**) Fluorescence emission of Alexa Fluor 647 in human serum and (**D**) in PBS. (**E**) Comparison of fluorescence emission of both human serum and PBS show no difference. (**F**) Signal to noise aspect ratios in human serum and in PBS. The signal-to-noise ratio is slightly reduced in human serum compared to PBS.

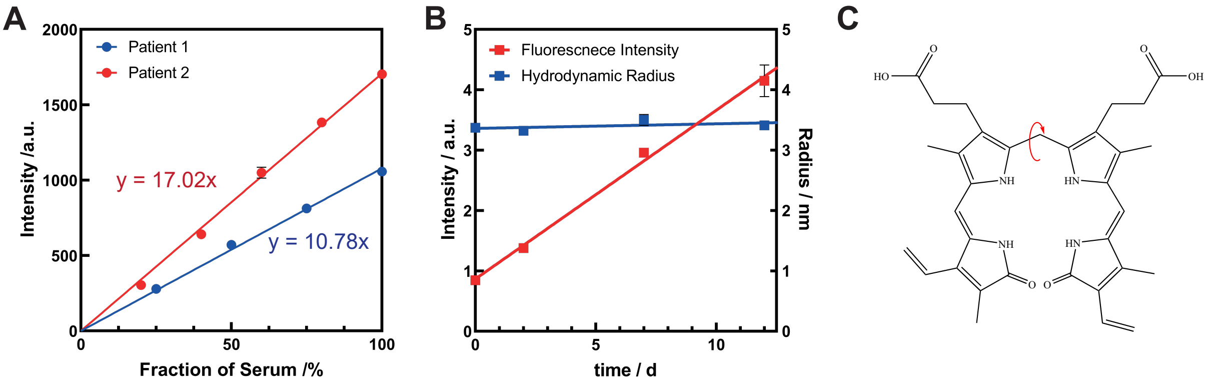


**Figure 3:** Fluorescence Emission of human serum, measured by microfluidic diffusional sizing. (**A**) Auto-fluorescence of human serum as a function of serum proportion for two different patients, showing that the behaviour varies among different individuals. (**B**) Increase in background intensity over time. As shown here, the background fluorescence increases linearly. The apparent hydrodynamic radius, $R_h$, remains unchanged. (**C**) Structure of bilirubin. The red arrow indicates the rotation which is hindered by complexation of bilirubin to HSA, a possible source of the fluorescence of the human serum.

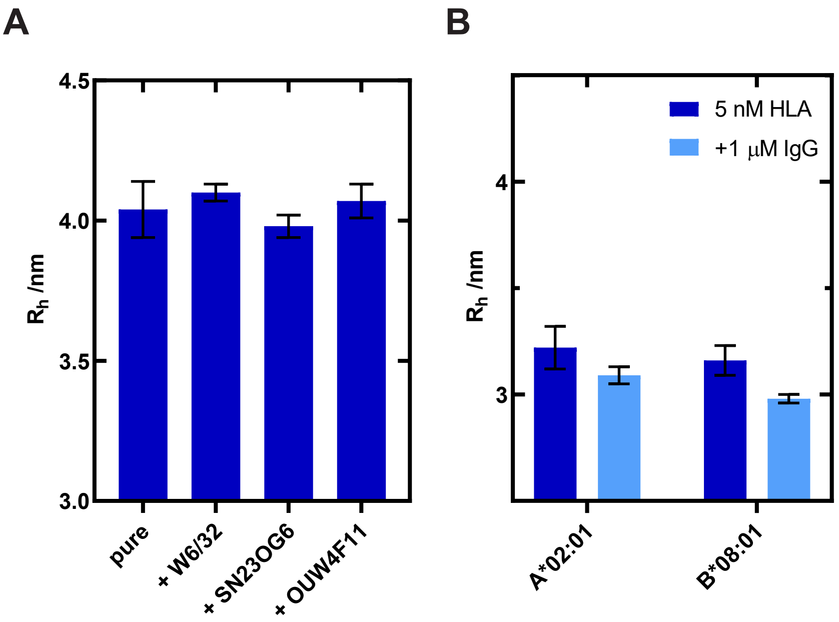


**Figure 4:** Control experiments testing specificity of binding interactions. (**A**) Comparison of hydrodynamic radii of Alexa Fluor 647 labelled BSA, both pure and after incubation with different antibodies. This demonstrates that the HLA specific antibodies do not recognise the fluorophore, thus, every binding interaction determined can be assumed to be specific. (**B**) Hydrodynamic radii of different HLA variants determined purely or after incubation with 200 nM IgG (ab205198), showing no size increase and, thus, suggesting selective interaction between these HLA variants and specific alloantibodies.

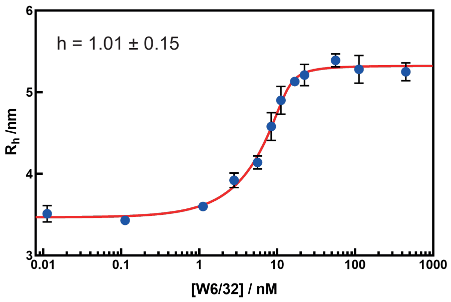


**Figure 5:** Binding curve of 25 nM HLA A*03:01 with varying concentration of antibody W6/32. The blue points are averages ofthree replicates, and the red line is the fit according to a Hill equation. From this data, the Hill coefficient $h = 1.01 \pm 0.15$ could be determined.

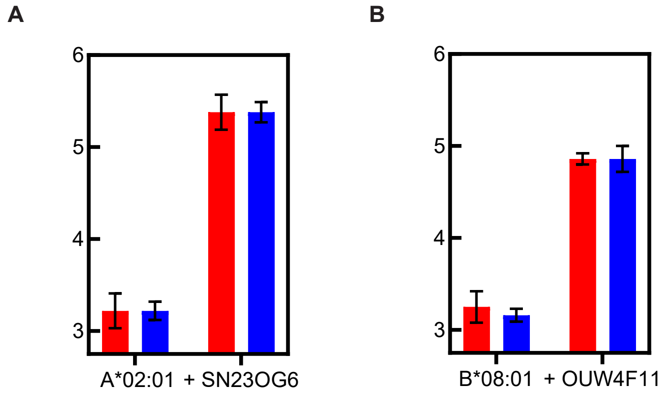


**Figure 6:** Hydroydnamic radii measured for (**A**) the interaction between 5 nM HLA A*02:01 and 1$\mu$M SN23OG6 and (**B**) the interaction between 5 nM HLA B*08:01 and 1$\mu$M OUW4F11, as shown in Fig. 3C-D. The data here, in comparison to the binding curves, were recorded on the same day, reducing batch-to-batch variability.

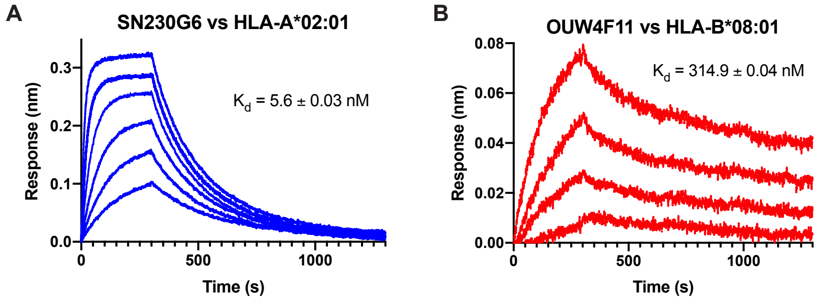


**Figure 7:** Biolayer Interferometry Curves for (**A**) the interaction between SN23OG6 vs. HLA A*02:01 ([HLA] = 100 nM, 50 nM, 25 nM, 12.5 nM, 6.25 nM and 3.125 nM) and (**B**) the interaction between OUW4F11 and HLA B*08:01 ([HLA] = 4000 nM, 2000 nM, 1000 nM, 500 nM).
